# Two polymorphs of an independent amyloid core in medin’s C-terminal region

**DOI:** 10.64898/2026.10.29.762820

**Authors:** Brajabandhu Pradhan, Frederic Rousseau, Joost Schymkowitz

## Abstract

Medin, a 50-residue fragment of lactadherin (MFG-E8), forms aortic medial amyloid, the most common localized amyloid in humans, and co-deposits with vascular amyloid-β (Aβ) in Alzheimer’s disease. In the full-length medin fibril the N-terminal half forms the entire protofilament interface, yet the strongest predicted amyloidogenic determinants lie in the C-terminal region. Here we show that this region aggregates on its own. The isolated C-terminal aggregation-prone region (medin C-APR, medin 31–53) begins to aggregate with the same lag time as full-length medin and assembles into two coexisting polymorphs, determined here by cryo-EM at 2.71 and 2.91 Å. Both are built from the same two-stranded β-arch and a pseudo-2₁ protofilament pair closed by an aromatic–aliphatic zipper, and differ only in how many accessory chains the exposed hydrophobic face recruits. The medin C-terminus therefore carries an amyloid core of its own, unrelated to that of the full-length fibril: medin has more than one aggregation-prone region, and the structural contribution of each region depends on sequence and assembly context.

## Introduction

Aortic medial amyloid (AMA) is the most common form of localized amyloid in humans. Its principal constituent, medin, is a 50-residue internal fragment of lactadherin/MFG-E8 (lactadherin residues 268–317) that is deposited in the medial layer of the upper thoracic aorta and other large arteries in virtually all individuals over the age of 60 ^1^. Unlike systemic amyloidoses, in which a destabilizing mutation or a proteostatic failure precipitates deposition, medin deposition is a near-universal correlate of vascular ageing. Medin deposits are found in the arteries of most older individuals, including the temporal arteries, where they occur both with and without giant-cell arteritis ^2^. They are implicated in the ageing loss of aortic elasticity ^3^ and in the pathogenesis of thoracic aortic aneurysm and dissection^4^. Medin aggregation causes cerebrovascular dysfunction in ageing wild-type mice ^5^, and vascular smooth-muscle-cell senescence accelerates medin aggregation through small extracellular vesicles ^6^. Medin co-aggregates with vascular amyloid-β (Aβ) in Alzheimer’s disease and promotes its deposition ^7^, linking vascular ageing with neurodegeneration. Medin is also among the fastest-aggregating amyloid peptides characterized, with a fibril elongation rate constant higher than those of amyloid-β, tau and α-synuclein ^8^.

Structural information on medin amyloid fibrils has remained limited. Solution NMR of the C2 domain of MFG-E8, from which medin is released, defined that domain’s fold and showed how its loops recognize phosphatidylserine ^9^. Molecular-dynamics and AlphaFold2 modelling of the whole protein then found that MFG-E8 favours a compact conformation in which the medin region, and its C-terminal cleavage site in particular, is poorly accessible to solvent, implying that the protein must unfold at least partly before medin can be excised ^10^. Both describe the folded precursor rather than the amyloid. Solid-state NMR of medin and medin-derived peptides established a β-rich architecture with disordered flanks and proposed a stabilizing Asp25–Lys30 salt bridge ^11–13^, and ¹³C-direct-detection NMR with structural bioinformatics probed the monomer and its nucleation ^14^. We recently determined cryo-EM structures from mixtures of medin and Aβ40, which included the first atomic structure of full-length medin fibrils ^15^. There all 50 residues are ordered: medin 1–26 occupies the interior of the protofilament and forms the entire interface between the two protofilaments, while the C-terminal region (Gly27–Ala50) runs along the outside of the fibril, exposed to solvent.

However, several studies have identified the medin C-terminus as an amyloidogenic region. Two sequence-based predictors place amyloid-competent segments in both halves of medin (Fig. 1a). WALTZ ^16^, run at its high-sensitivity threshold, calls medin 4–27 (Lys4–Gly27; peak score 94.6 at Lys4) and 32–50 (Val32–Ala50; peak 90.3 at Val32); at its default threshold only the N-terminal segment is called. CORDAX ^17^, which scores the structural compatibility of a sequence with an amyloid core, predicts four segments: medin 7–13 (Asn7–Ala13), 17–27 (Gly17–Gly27), 31–39 (Glu31–Gly39) and 44–49 (Gly44–Val49), the last two separated by a gap at Ala40–Phe43. The highest-scoring segment anywhere in medin is 31–39, where residues Gly34–Gly39 all reach a score of 0.86, above either N-terminal segment. Both predictors thus identify the C-terminal region as amyloid-competent. Solid-state NMR shows that medin fibrils are predominantly β-sheet with disordered elements, and that full-length medin and the C-terminal fragments medin 30–50 and medin 42–49 all aggregate, although they pack differently, implying that at least two of the three predicted amyloidogenic regions are needed to build a medin fibre ^11,12^. This underscores the importance of the medin C-terminal aggregation-prone region (C-APR) in amyloid formation. Here we asked whether the medin C-APR can act independently, and how the fibrils it forms compare with the full-length medin fibril assembled in the presence of Aβ40.

**Fig. 1.**
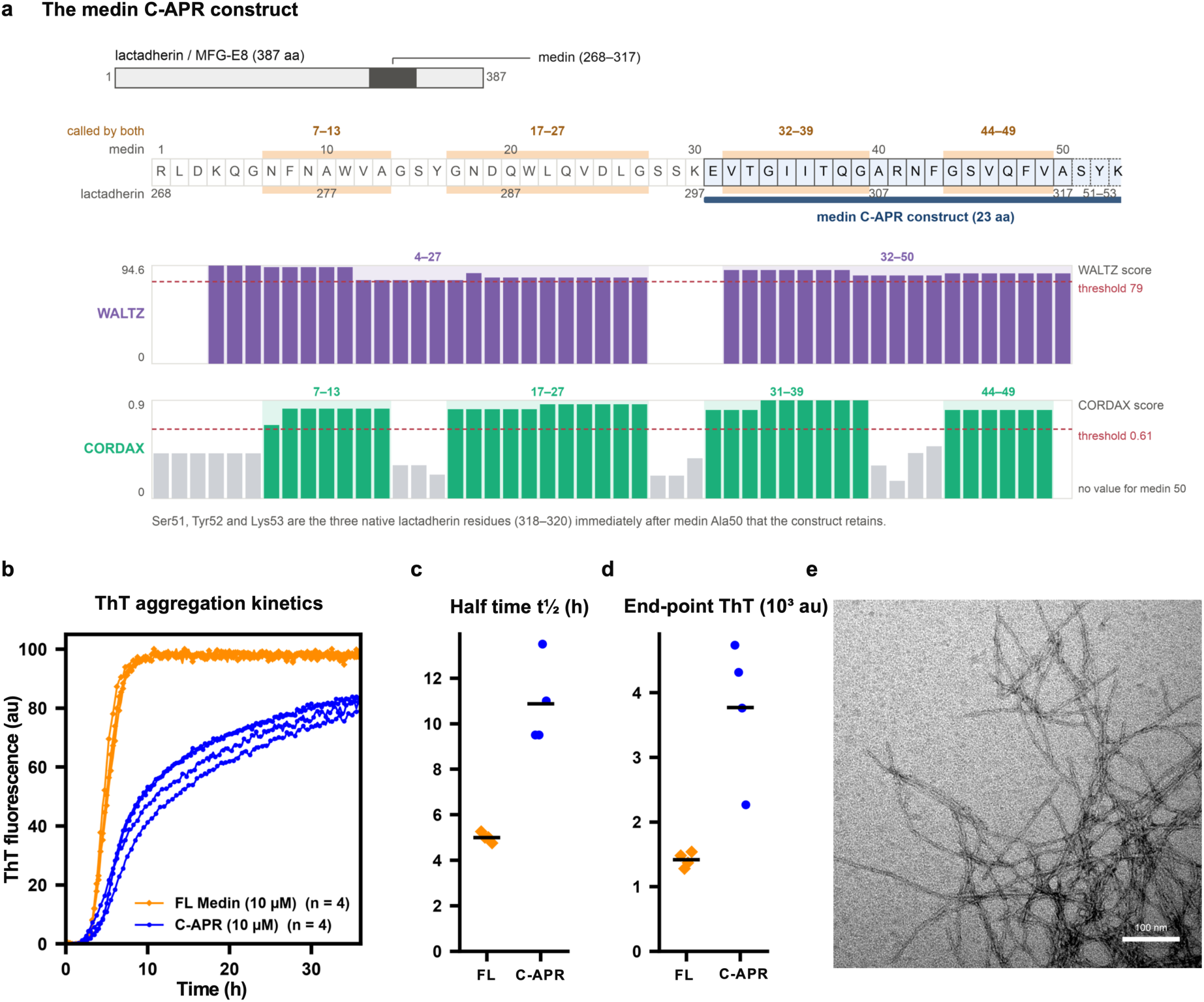
The medin C-APR is an aggregation-prone region that forms fibrils on its own. **a**, The medin sequence in its lactadherin context, with dual numbering (medin 1–50 = lactadherin 268–317) and the synthetic C-APR construct bracketed; tracks below give the per-residue aggregation propensity predicted by WALTZ and CORDAX, with shading where both predictors agree. b, ThT fluorescence of the medin C-APR and of full-length medin, both at 10 µM in PBS (pH 7.4) with 10 µM ThT at 37 °C; individual wells (n = 4 per peptide), each normalized to its own maximum. c, Half-time t½ of each well, taken as the first crossing of the half-maximum of that trace; bars, means. d, End-point ThT signal of each well, the median of the last 10 reads minus the starting level; bars, means. e, Negative-stain TEM of C-APR fibrils grown quiescently at 100 µM, the preparation used for cryo-EM; 640 nm field, scale bar 100 nm.

To test this, we studied the structures of fibrils formed by the isolated medin C-APR. The construct is lactadherin 298–320, which runs from medin Glu31 to Ala50 and on into the three following lactadherin residues. We number these as a continuation of medin, Ser51 to Lys53, so that every residue carries one number; the deposited models keep lactadherin numbering, 267 higher throughout. The same peptide, there called CT-medin, has been studied before: Wagner et al. synthesized it as the medin species they most consistently identified in tissue extracts by mass spectrometry ^7^. We show by thioflavin-T (ThT) kinetics that the C-APR starts to aggregate as readily as full-length medin, and we determine cryo-EM structures of two coexisting C-APR polymorphs. Both are built from the same V-shaped subunit, a two-stranded β-arch closed onto a dry aromatic zipper. That subunit has no counterpart in the full-length fibril core, establishing the medin C-terminus as a second, independent amyloid core.

## Results

### The medin C-APR aggregates on its own, with a slower growth phase than full-length medin

To test whether the C-APR reproduces the aggregation of full-length medin, we monitored its aggregation kinetics by ThT fluorescence (Fig. 1b). Both peptides were followed at 10 µM in PBS (pH 7.4) at 37 °C, with a brief shake before each read. The medin C-APR showed a lag time of 3.6 ± 0.6 h and a half-time t½ = 10.6 ± 2.0 h, compared with 3.11 ± 0.07 h and 4.9 ± 0.3 h for full-length medin (n = 4 wells per peptide, Fig. 1c). The maximum growth rate, taken from the steepest part of each normalized trace, was 0.11 ± 0.03 h⁻¹ for the C-APR and 0.32 ± 0.05 h⁻¹ for full-length medin. The end-point ThT signal was about 2.7-fold higher for the C-APR (3.8 ± 1.1 × 10³ au) than for full-length medin (1.42 ± 0.12 × 10³ au; Fig. 1d); because ThT fluorescence per unit of fibril can differ between fibril types, we do not read this as a difference in the amount of fibril formed. The two constructs therefore begin to aggregate at the same time, and differ in what follows: the C-APR takes about twice as long to reach half-completion. The C-APR therefore initiates aggregation with similar kinetics to full-length medin but grows more slowly.

The fibrils used for cryo-EM were grown separately, quiescently at 100 µM for 9 days and without dye (Methods), and they differ in morphology from the fibrils that the aggregation assay produced. The ThT assay resulted in a heterogeneous mixture of short rods, bundles and thinner isolated fibrils (Supplementary Fig. 1c), and was not suitable for helical reconstruction. The quiescent preparation reproducibly gave long, unbranched fibrils (Fig. 1e), in our two independent preparations (Supplementary Fig. 1a,b). Full-length medin at the end of the assay formed long, unbranched fibrils (Supplementary Fig. 1d). The kinetics above therefore report how readily each peptide aggregates, while the structures below describe the quiescently grown C-APR fibril.

### Cryo-EM resolves two coexisting medin C-APR polymorphs

Fibrils grown quiescently at 100 µM were vitrified and imaged by cryo-EM (Methods; Supplementary Fig. 3). Two-dimensional classification of segments from 1,501 curated micrographs, in CryoSPARC v5^18^, separated two fibril morphologies. The two morphologies coexist side by side as 87 % of the micrographs contained both morphologies (Fig. 2a,b). Morphology 1 is the more abundant, with 65 % of the assigned segments and Morphology 2 the remaining 35 % (Fig. 2d). Helical reconstruction with C1 point-group symmetry gave maps at 2.71 Å for Morphology 1 and 2.91 Å for Morphology 2 (gold-standard FSC = 0.143; Fig. 2c; Supplementary Fig. 2). Morphology 1 refined to a twist of −2.75° and a rise of 4.753 Å while Morphology 2 to −2.37° and 4.77 Å, giving crossover distances of about 311 and 362 Å respectively. We built atomic models de novo into both maps (Figs. 3 and 4). Fig. 2e summarizes the residues ordered in each chain, Supplementary Fig. 4 the map–model fit, and Supplementary Table 1 the refinement and validation statistics.

**Fig. 2.**
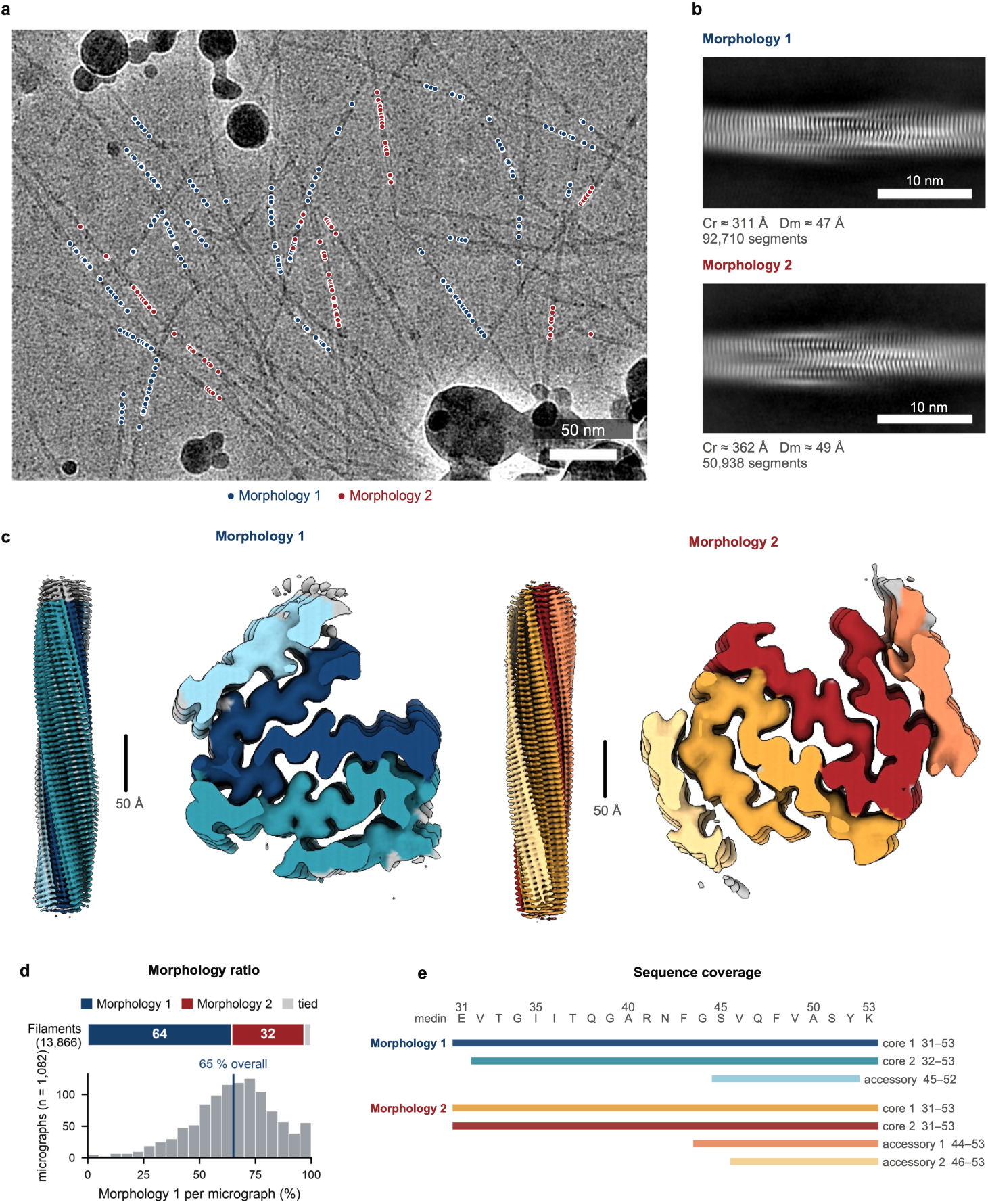
Cryo-EM of medin C-APR fibrils reveals two coexisting morphologies. **a**, Motion-corrected micrograph (0.831 Å per pixel, low-pass filtered to 8 Å) with the traced filaments coloured by morphology: Morphology 1 (dark blue) and Morphology 2 (dark red), Scale bar, 50 nm. b, Reference-free 2D class averages of each morphology (filament axis horizontal) with the crossover distance (Cr, from the refined helical parameters), the width (Dm, full width at 25 % of the class projection profile), segment numbers. Scale bar, 10 nm. c, Sharpened reconstructions of Morphology 1 (2.71 Å) and Morphology 2 (2.91 Å), side view and cross-section, coloured by chain type (Morphology 1 in blues, Morphology 2 in warm tones; the protofilaments in the deeper shade and each accessory segment in a lighter tint of the protofilament it caps); both morphologies are shown at the same scale. Scale bar, 50 Å. d, Relative abundance of the two morphologies in this preparation. Top, traced filaments with at least three assigned segments, assigned by majority vote (8,925 Morphology 1, 4,506 Morphology 2, 435 tied, grey); the underlying segment counts are 89,563 and 47,791, excluding the 3,147 segments common to both refinements. Both morphologies were present on 87 % of the 1,420 micrographs that contributed segments. e, Residues ordered in each chain of one layer, in medin numbering. Both cores span essentially the whole construct, whereas the accessory segments are confined to its C-terminal third.

**Fig. 3.**
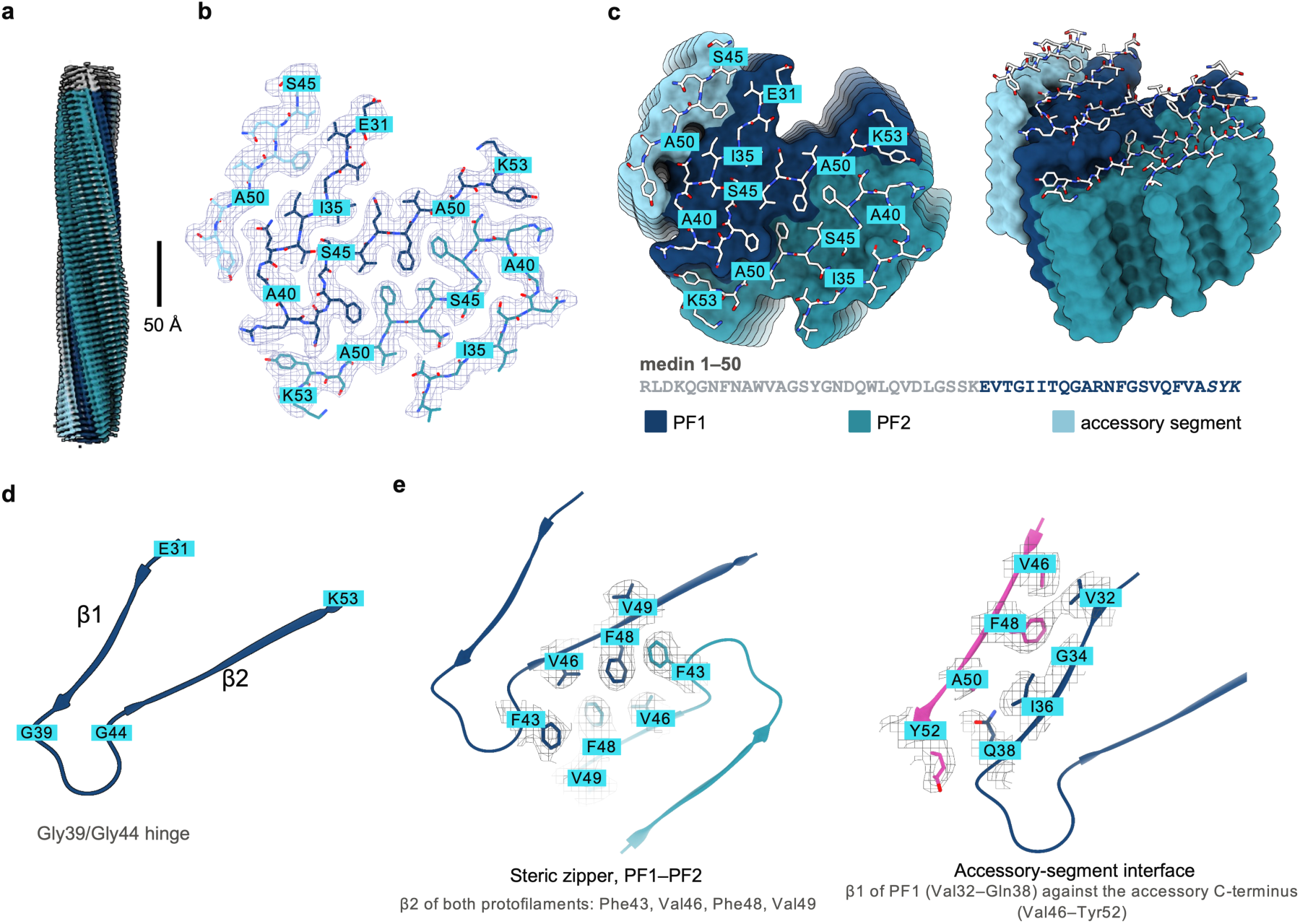
Architecture of Morphology 1. **a**, The sharpened map along the fibril axis, coloured by chain (PF1 dark blue, PF2 teal, accessory segment pale blue). Scale bar, 50 Å. b, One cross-β layer as mesh with the model, labelled every fifth residue. c, Surface of a six-layer stack seen down the fibril axis and tilted, with the top layer as sticks; below, the C-APR construct within the medin sequence. d, One protofilament subunit as a cartoon: β1 Val32–Gln38, turn Gly39–Gly44, β2 Ser45–Lys53, with the glycines that bracket the turn marked. e, The two interfaces in close-up, transverse (down the axis, one layer) and lateral (layers i, i−1, i−2): left, the Phe43/Val46/Phe48/Val49 zipper between PF1 and PF2; right, the accessory segment against the β1 face of PF1.

**Fig. 4.**
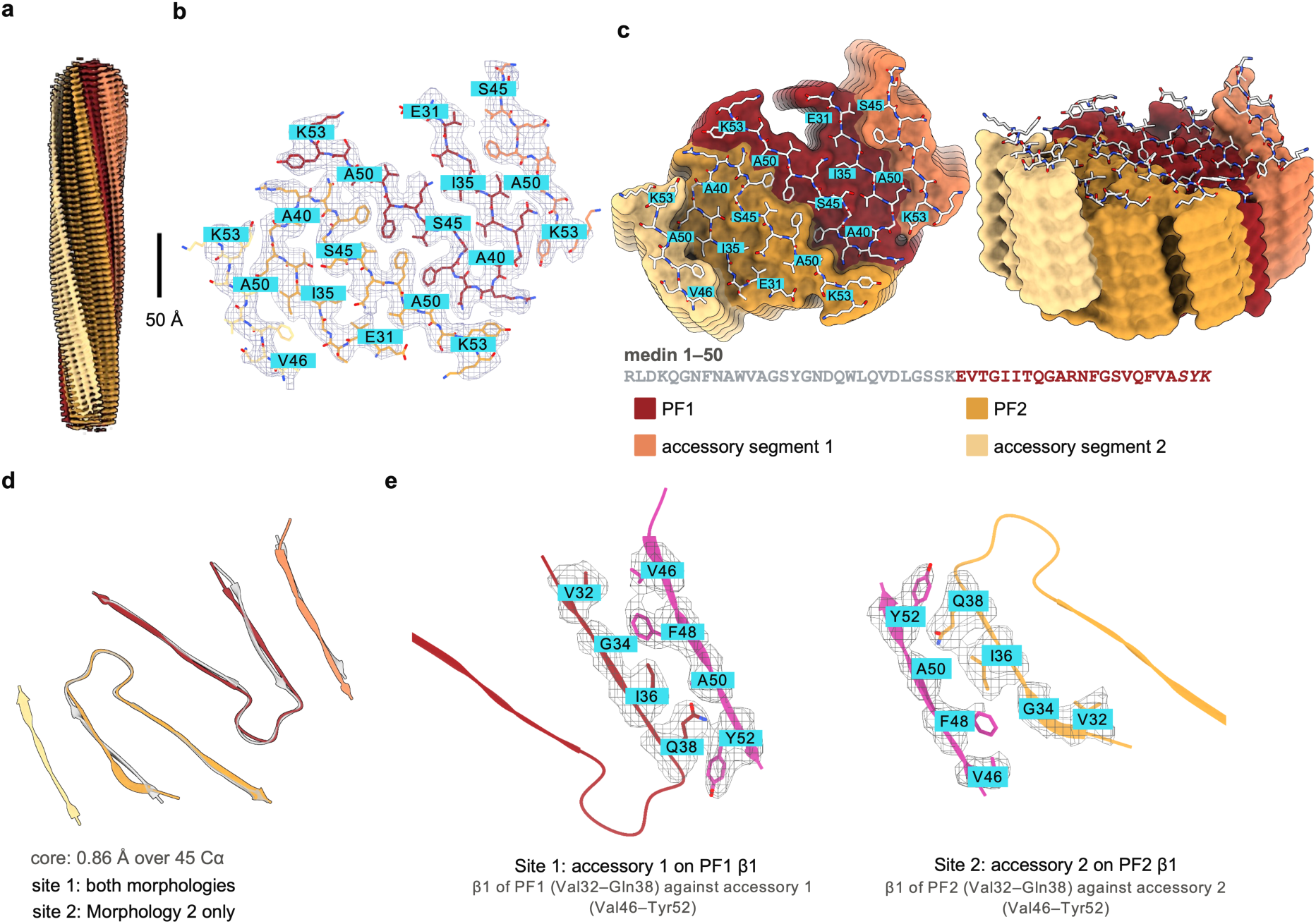
Architecture of Morphology 2. Panels a–c as in Fig. 3, for the Morphology 2 map and model: a, the map along the fibril axis, coloured by chain (PF1 dark red, PF2 gold, accessory segments 1 and 2 salmon and pale gold); b, one cross-β layer as mesh with the model; c, the six-layer surface, with the helical parameters and the construct sequence below. d, Morphology 1 (grey) superposed on Morphology 2 through PF1 and PF2 (0.86 Å over 45 Cα): the core and accessory site 1 are shared between the polymorphs, site 2 occurs only in Morphology 2. e, The two accessory sites in close-up, transverse and lateral: site 1, accessory segment 1 on the β1 face of PF1, where the Gln38 side chain contacts the segment’s main chain; site 2, accessory segment 2 on the β1 face of PF2, the symmetry-related site.

### Morphology 1: a pseudo-2₁ protofilament pair coated by one accessory segment

Each cross-β layer of Morphology 1 contains three medin C-APR chains resolved to different extents (Fig. 3). The core of the fibril is composed of two C-APR chains: protofilament 1 (PF1), traced from Glu31 to Lys53, and protofilament 2 (PF2), from Val32 to Lys53, which is missing one N-terminal residue relative to PF1. A third, accessory chain coats the outer surface of PF1 and is ordered only over its C-terminus, spanning Ser45–Tyr52; its N-terminal two-thirds are not resolved, likely due to their flexibility. Thus, the two core chains are largely ordered, whereas the peripheral accessory chain is only partially resolved.

A single protofilament subunit folds into a two-stranded topology (Fig. 3d): β1 spans Val32–Gln38 and β2 spans Ser45–Lys53, connected by the six-residue segment Gly39–Gly44. The chain makes a ∼180° turn at Asn42 in PF1 and Phe43 in PF2, where the backbone adopts a left-handed conformation that facilitates the reversal. The two glycines bracketing this segment introduce a 90° kink in the connector. The loop corresponds to a region of low predicted aggregation propensity in CORDAX, whereas both β-strands coincide with regions of high predicted propensity.

PF1 and PF2 are related by an approximate two-fold screw (pseudo-2₁) axis. Superposition of the two subunits gives an RMSD of 1.41 Å, with the largest differences localized to the N-terminus, at Val32 and Thr33; this difference may reflect the absence of an additional accessory chain at the periphery of PF2. The interface between the two subunits is formed by the β2 strands, which likely explains why the RMSD is lowest (a mean of 0.8 Å) across this region. In each subunit, the two strands sit roughly 14 Å apart, forming an open arch rather than a self-contained steric zipper, with intramolecular side-chain contacts confined to the closure point, where Thr33, Ile35, and Thr37 pack against Ser45 and Gln47.

The PF1–PF2 interface is a tight contact formed by hydrophobic and aromatic residues. Here, Val46, Phe48 and Val49 of PF1 pack against Phe43, Val46 and Phe48 of PF2. The phenylalanines cross-pair across the interface, Phe48 against Phe43, and stack along the fibril axis (Supplementary Fig. 5a). At the periphery, Arg41 and Asn42 of one protofilament contact Tyr52, Ser51 and Ala50 of the other, and the pseudo-2₁ symmetry repeats the same contacts in reverse. Since every one of these contacts is apolar or aromatic, and only one nitrogen-oxygen pair across the interface is close enough to hydrogen-bond, the protofilaments meet through a dry, desolvated zipper rather than a water-mediated polar contact.

The exposed β1 face of protofilament 1 interfaces with an accessory chain, ordered only over its hydrophobic C-terminal stretch. The accessory chain sits between two layers of β-strands (Supplementary Fig. 5b) and packs its own C-terminal Phe48, Val46 and Ala50 against the outer face of PF1’s β1 strand (Val32, Gly34, Ile36 and Gln38). In this way, the accessory segment occupies the partly unsatisfied hydrophobic face of the PF1 β1 strand and also makes its N-terminal residue ordered.

### Morphology 2 shares the core but recruits two accessory segments

Morphology 2 contains four chains per layer (Fig. 4): two protofilament subunits, PF1 and PF2, both traced from Glu31 to Lys53, and two accessory segments ordered over Gly44–Lys53 and Val46–Lys53. As in Morphology 1, PF1 and PF2 are related by an approximate two-fold screw. Superposing the Morphology 2 core on that of Morphology 1 gives an r.m.s.d. of 0.86 Å over 45 Cα atoms, so the β-arch and the zipper are conserved between the polymorphs. The zipper at the interface between the protofilaments is again built from Phe43, Val46, Phe48 and Val49, and makes no cross-interface hydrogen bond at all. The difference between the two morphologies lies in the accessory chains: here two segments cap the surface instead of one.

The two segments occupy equivalent sites on PF1 and PF2. Accessory segment 1 binds the β1 face of PF1 at the position used by the Morphology 1 accessory chain, with its Phe48 against Ile36 and Gly34 of the next layer. Accessory segment 2 binds the β1 face of PF2 through Tyr52 and Phe48. The pseudo-2₁ axis that relates PF1 and PF2 also maps one accessory segment onto the other, to within 1.7 Å. Each accessory chain lies about 0.6 of a rise from the protofilament it caps, one above and one below (Supplementary Fig. 5c,d). The two accessory chains differ in how much is ordered: Gly44–Lys53 is ordered for the accessory chain of PF1, while Val46–Lys53 is ordered for that of PF2, and correspondingly in the extent of contact. The Morphology 2 cross-section is correspondingly wider, 57.9 Å against 47.2 Å for Morphology 1. The three ionizable side chains of the construct, Glu31, Arg41 and Lys53, all point into solvent in both polymorphs, and neither structure contains a buried salt bridge or charge pair.

We then compared the C-APR structures with the full-length medin fibril, grown in co-aggregation with amyloid-β40 ^15^. The two structures use different parts of the sequence to build their cores (Supplementary Fig. 6). In the full-length medin fibril all 50 residues are ordered, but the two protofilaments pack exclusively through the N-terminal half: every contact across the protofilament interface comes from medin 1–30, and within that from residues 1–4 and 21–23 alone. Medin 31–50 touches the opposing protofilament nowhere; it is held only by stacking within its own protofilament, running along the fibril surface as an extended arm 63 Å from end to end. In both C-APR fibrils the same twenty residues instead fold back on themselves into a compact β-arch about 15 Å from end to end and form the entire core. Superposing the shared segment gives a backbone r.m.s.d. of 16.5 Å. The C-terminal region therefore adopts substantially different conformations in the two fibril structures. Apart from that, the previous study also had Aβ40 present in solution, whose role in shaping the structure cannot be determined.

### Energetics of the C-APR fibrils in the context of known amyloid structures

To place the C-APR folds among known amyloid structures, we computed per-residue FoldX stabilities for every protofilament stack and compared them with the 1,185 protofilament stacks in the Amyloid Explorer database (Fig. 5 and Supplementary Fig. 7). We follow the Amyloid Explorer convention throughout: a residue with ΔG_contrib below zero stabilizes the fold, one above zero is structurally frustrated. The most stabilizing residues of both polymorphs lie in the zipper and at the closure of the arch: Phe48, Val46, Phe43 and Ile36 each have ΔG_contrib between −0.9 and −2.8 kcal mol⁻¹. The most frustrated are Arg41 (+2.0 to +2.5 kcal mol⁻¹) and the turn glycines (+0.6 to +2.4) in both, together with the terminal lysine wherever it is ordered (+1.7 to +6.1) (Fig. 5a). The core stacks have mean residue ΔG values of −0.04 to +0.40 kcal mol⁻¹, spanning the 28th to 66th percentile of Amyloid Explorer protofilament stacks, so both folds sit within the range of experimentally observed amyloid cores (Supplementary Fig. 7a). The accessory segments behave quite differently in the two polymorphs. In Morphology 1 the single segment is as favourable as the cores it caps, −0.01 kcal mol⁻¹ per residue against −0.04 and +0.40, the 29th percentile of the database against their 28th and 66th. In Morphology 2 the two accesory segments differ from one another: one is unremarkable at +0.10 kcal mol⁻¹ per residue, the 40th percentile, while the other is the most frustrated stack of either structure at +0.73, the 86th percentile. This is consistent with their peripheral location and weaker density. About half the core residues of each polymorph are frustrated, which falls within the range observed for other amyloid cores, against two-thirds or more of the accessory segments.

**Fig. 5.**
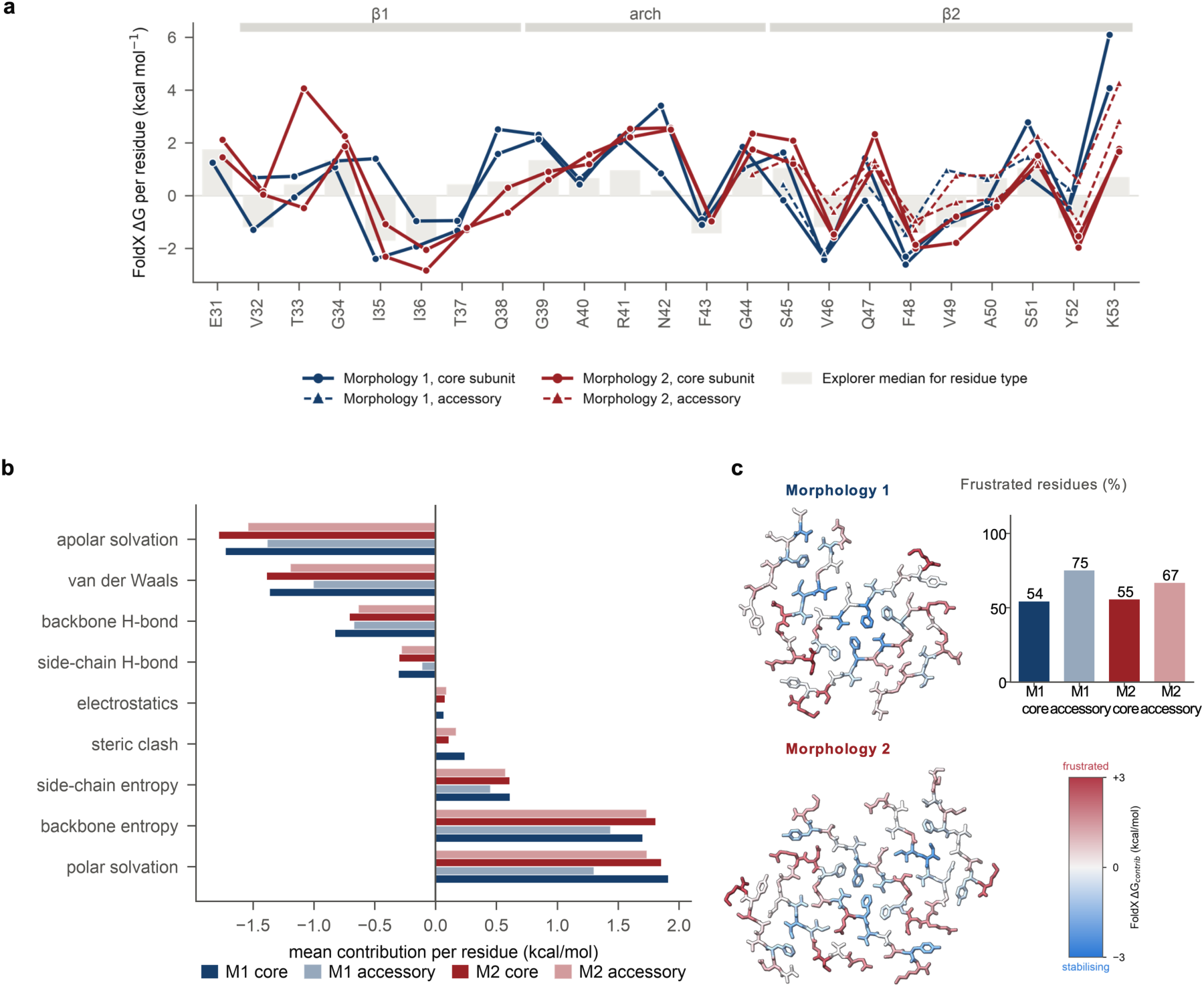
Energetics of the medin C-APR fibrils. **a**, Per-residue FoldX ΔG per protofilament stack (circles, core; triangles, accessory). b, Mean per-residue contribution of each FoldX term for the core and accessory stacks of both morphologies. c, One cross-β layer of each morphology coloured by per-residue ΔG_contrib on a scale symmetric about zero, both at one orthographic scale, with the proportion of residues that are structurally frustrated (ΔG_contrib > 0) in each stack (inset).

Decomposition of ΔG_contrib into its physical terms identifies the interactions that stabilize each fold and the costs they offset (Fig. 5b). Stabilization in both polymorphs derives from apolar solvation, about −1.8 kcal mol⁻¹ per core residue, and van der Waals packing, about −1.4, with backbone hydrogen bonding contributing less than half as much; polar solvation (about +1.9) and the backbone entropy lost on ordering (+1.7 to +1.8) oppose them. The accessory segments show the same profile with the large terms attenuated, consistent with their smaller buried surface. Steric clash is the exception: it is negligible in the single accessory segment of Morphology 1, at +0.01 kcal mol⁻¹ per residue, but reaches +0.11 to +0.24 in every other stack, so that segment docks without strain while the two of Morphology 2 do not. Projecting ΔG_contrib onto one cross-β layer shows the same distribution in space (Fig. 5c): frustration increases with distance from the fibril axis, the innermost third of residues being favourable on average and the outermost third not. The trend is not a strict partition, since a minority of buried residues is frustrated as well, and the most frustrated positions are the turn glycines, Arg41 and the ordered C-terminal residues.

The same calculation on the full-length medin fibril shows where the two folds differ energetically (Supplementary Fig. 6d). Over medin 31–50, the stretch the two constructs share, the segment is similarly favourable in the two contexts. The difference is confined to the connector. In the full-length fibril, where these residues run as a single extended strand, Ala40–Phe43 carries +0.39 kcal mol⁻¹ per residue; in the C-APR fibrils, where the same residues close the β-arch, it carries +0.94 in Morphology 1 and +1.33 in Morphology 2. The turn is thus two to three times as frustrated in both C-APR folds as in the full-length fibril, so its cost, rather than anything about the strands, is the energetic feature both C-APR folds share and the full-length fibril lacks. Asn42 (+0.24 in the full-length fibril against +2.33 in the C-APR cores) and Ser45 (+0.25 against +1.19) carry most of it.

## Discussion

The structures reported here show that the medin C-APR, on its own, forms an ordered amyloid core that is distinct from the core of the full-length medin fibril.

First, medin contains more than one amyloid-competent region. The isolated C-APR builds a complete amyloid core from residues that, in the full-length fibril, lie outside the protofilament interface ^15^, and closes it with a two-stranded β-arch and a dry zipper. Because the C-APR starts to aggregate as readily as full-length medin, with the same lag time, the C-terminal region can independently support amyloid formation, even though it lies in an extended form and exposed to the solvent in the full-length fibril.

Second, the C-APR structures clarify the relationship between predicted aggregation-prone segments and the observed fibril architecture. The segments flagged by the predictors are not competing nuclei but the two strands of one β-arch joined by a glycine-bracketed turn. The CORDAX predictions correspond closely to the observed secondary-structure elements, as it assigns the highest propensity to residues 31–39 and 44–49 and leaves a gap at Ala40–Phe43, which is exactly the turn that connects β1 and β2 in both polymorphs. Therefore, the predicted hotspots and the observed strands coincide residue for residue.

Third, both polymorphs leave one β1 face of the core exposed and cap it with an additional chain. Morphology 1 recruits one accessory segment and Morphology 2 two, at the two sites related by the pseudo-2₁ axis. The comparison with the full-length fibril suggests why. Stripped of its accessory chain, the C-APR core leaves 150 Å² of apolar surface on the β1 face; the accessory buries most of it, docking through its own Val46, Phe48 and Ala50. The same face is not exposed in the full-length fibril, where it is already covered by residues of the N-terminal half. In the full-length fibril, this hydrophobic surface is buried by intramolecular contacts, whereas in the C-APR fibril it is covered by an additional peptide chain. It also offers an explanation for why the accessory chains are only partly ordered. The synthetic peptide is a single species, so they are most likely complete C-APR molecules whose N-terminal two-thirds remains disordered, because the core offers a site for the short hydrophobic C-terminal stretch and none for the rest. Consistently, they carry the highest proportion of frustrated residues in either structure, and the single most frustrated stack of all is one of the two in Morphology 2 (Fig. 5b and Supplementary Fig. 7a), as expected of subunits held by burial alone. The C-APR core carries a second, unrelated cost. It leaves Arg41 fully exposed in both polymorphs, with no acidic group anywhere near it, whereas the same residue is buried in the full-length fibril, coordinated by backbone and amide oxygens of medin 14–18. Stacked every 4.75 Å, an exposed Arg41 becomes a ladder of like charges along the fibril axis, which is why FoldX scores it among the most destabilizing core residues of every core stack in both polymorphs. Compared with the full-length fibril, the C-APR core lacks both the intramolecular contacts that shield its hydrophobic surface and the interactions that stabilize Arg41. The former are partly replaced by accessory-chain binding, whereas Arg41 remains solvent-exposed. One caveat: the full-length fibrils were assembled in the presence of amyloid-β, so the absence of the N-terminal half and the absence of amyloid-β are confounded here. The structures show that the N-terminal half can perform this capping role, not that its absence caused the recruitment. A core that recruits extra chains through the same exposed hydrophobic face is the kind of template that should support heterotypic interactions, as is known for medin with Aβ ^7,15^.

Fourth, FoldX analysis indicates that hydrophobic packing and van der Waals interactions are the principal stabilizing contributions in both polymorphs, whereas energetically unfavourable residues are more frequent at the fibril periphery. Apolar solvation and van der Waals packing carry both cores against polar solvation and backbone entropy, and energetically unfavourable residues lie preferentially, though not exclusively, towards the solvent-exposed surface (Fig. 5b,c). That the accessory segments of Morphology 2 are slightly more strained than the cores they bind suggests the second recruitment site is accommodated rather than optimal. This may help explain the coexistence of the two polymorphs in one preparation: the core is the same in both, and what distinguishes them is the extent of accessory-chain binding to the exposed β1 surface. If that is right, the polymorph ratio should be sensitive to the conditions that change the supply of free peptide, a prediction that a concentration series, which we have not yet run, would test directly.

Despite its links to cerebrovascular dysfunction and to vascular Aβ deposition in Alzheimer’s disease, medin remains among the least characterized amyloids structurally. Showing that its C-terminal region carries an amyloid core of its own, unrelated to the core of the full-length fibril, means that medin can adopt structurally distinct amyloid cores. For tau, TDP-43 and α-synuclein, particular folds are now tied to particular diseases ^19–21^, and that correspondence is what gives a structure its diagnostic weight. No such correspondence can yet be drawn for medin. How medin is released from lactadherin in vivo is also unresolved, and our results suggest that where the protein is cut may decide which core it builds: full-length medin and the C-APR begin to aggregate with the same lag time, yet assemble into entirely different folds. Several limitations remain. The comparison with the full-length fibril is confounded three ways at once, a different construct, a different buffer and the presence of Aβ40, so no single factor can be assigned as the cause of the different fold, though the coexistence of two polymorphs in one preparation shows that polymorphism here arises with no difference in environment at all. The kinetics and the structures were measured on separately grown material, the assay end point being heterogeneous and unsuitable for cryo-EM, so the kinetics speak to aggregation propensity rather than to the polymorphs described here. Finally, both polymorphs were obtained in vitro. Whether either structure occurs in human aortic medial amyloid remains unknown and will require structural analysis of patient-derived material.

## Methods

### Peptide production

The medin C-APR (lactadherin residues 298–320; EVTGIITQGARNFGSVQFVASYK, corresponding to medin Glu31–Ala50 followed by lactadherin Ser318–Lys320, numbered here Ser51–Lys53) and full-length medin (residues 1–50) were synthesized in-house by solid-phase peptide synthesis as described previously ^22^ and stored as lyophilized powder (97.5 % by RP-HPLC; identity confirmed by LC–MS, Supplementary Fig. 8). To remove pre-formed aggregates, peptides were dissolved in 100 % 1,1,1,3,3,3-hexafluoro-2-propanol (HFIP) in glass vials, incubated for 10 min at room temperature and dried under a stream of nitrogen to a thin film, as described previously ^15^. Films were redissolved in Milli-Q water immediately before use, and the stock solutions were passed through 0.22 µm Spin-X centrifuge filters (Costar 8160).

### Fibril assembly

Fibrils for structural analysis were grown from monomeric C-APR at 100 µM in PBS (pH 7.4) at 37 °C for 9 days without shaking and were collected once aggregation had reached completion. These cultures contained no ThT and were not shaken, so they were harvested on a fixed schedule rather than on the plateau of the ThT plates. Fibrils were taken directly from the growth tube without centrifugation or buffer exchange.

### Thioflavin-T aggregation kinetics

Monomeric stocks of the medin C-APR and of full-length medin were diluted in PBS (pH 7.4) containing 10 µM ThT to a final peptide concentration of 10 µM, and 100 µl per well were dispensed into µClear medium-binding half-area 96-well plates (Greiner 675096), which were sealed to limit evaporation. Wells containing buffer and ThT without peptide were included on every plate as a baseline control. Fluorescence was recorded at 37 °C in a FLUOstar Omega plate reader (BMG Labtech; 448/10 nm excitation, 482/10 nm emission), every 15 min for 118 h, with a brief shake before each read. The plate format, the dye and the brief shaking before each read differ from the quiescent, dye-free conditions under which the fibrils analysed structurally were grown, and the two experiments are reported separately. Each curve was baseline-subtracted. Lag times, half-times and growth rates were taken from the curve normalized to its own plateau, and the end-point signal from the unnormalized trace, as the median of the last 10 reads minus the starting level. Half-time (t½) was taken as the first crossing of the half-maximum of each trace. The growth rate was taken as the steepest slope of the normalized trace, from a five-point smoothed derivative, and the lag time (t_lag) as the intercept of that tangent with the baseline. The medin C-APR and full-length medin were run at the same concentration on the same plate. Data are means ± s.d. of 4 wells per peptide on one plate.

### Negative-stain TEM

Fibrils were taken directly from the quiescent growth tube, without centrifugation or buffer exchange. Aliquots (5 µl) were applied to glow-discharged Formvar/carbon 400-mesh copper grids (Agar Scientific AGS162-4) for 3 min. Excess liquid was blotted with Whatman no. 1 paper, and grids were washed with three 20 µl drops of Milli-Q water, stained with 2 % (w/v) uranyl acetate for 1 min, blotted and air-dried ^15^. Grids were imaged at 80 kV on a JEM-1400 transmission electron microscope (JEOL) with a LaB₆ filament, using an EMSIS Quemesa camera, at nominal magnifications between 8,000× and 50,000×; the pixel size was calibrated from the microscope scale bar and is 0.257 nm at 50,000×. The material shown in Fig. 1e is the 100 µM, 9-day quiescent preparation from which the cryo-EM grid was also made. A second session compared three grids on the same day: that quiescent preparation repeated after four months, full-length medin, and the end point of the thioflavin-T assay (Supplementary Fig. 1).

### Cryo-EM sample preparation and data acquisition

Holey-carbon grids (Quantifoil R1.2/1.3 Cu 300 mesh) were glow-discharged for 60 s at 5 mA (ELMO, Cordouan Technologies). Fibril suspension (3 µl) was applied in a Leica EM GP2 plunger at 22 °C and 98 % relative humidity, incubated for 1 min, blotted from the front side and plunge-frozen in liquid ethane ^15^. Data were collected at the electron Bio-Imaging Centre (eBIC), Diamond Light Source (UK), on a Titan Krios (Thermo Fisher Scientific; instrument ID 3415) operated at 300 kV with a BioQuantum energy filter (20 eV slit) and a K3 direct electron detector (Gatan) in counting mode (5,760 × 4,092 px), using EPU 3.11 (C2 aperture 50 µm, no objective aperture). A total of 4,905 movies were recorded at a nominal magnification of 105,000×, corresponding to a calibrated pixel size of 0.831 Å. Each movie had a 1.61 s exposure fractionated into 40 frames with a total fluence of ≈40 e⁻ Å⁻² (1.0 e⁻ Å⁻² per frame), and the applied defocus cycled from −1.0 to −3.0 µm in 0.3 µm steps.

### Helical reconstruction

Processing used CryoSPARC v5.0.6^18^. Movies were motion-corrected (Patch Motion Correction, K3 gain reference) and CTF parameters estimated (Patch CTF). Micrographs that contained fibrils and had a CTF fit of 6 Å or better were retained (1,501 of 4,905), filaments were traced with Filament Tracer, and segments were extracted with an inter-box distance of 21 Å. Iterative 2D classification separated the two morphologies. Initial twists were derived from crossover distances measured in 2D class averages. For each morphology an initial model was generated ab initio inside Helical Refinement from 5,000 randomly selected segments, starting from a 20 Å reference and with no input volume; the final refinement was then started from the map of the preceding refinement in the same series. Final reconstructions used Helical Refinement with non-uniform regularization ^23^ and global and local CTF refinement (refined Cs 2.50 mm) in a 400-pixel box (332.4 Å), C1 point-group symmetry and refined helical parameters (Morphology 1: −2.75°, 4.753 Å; Morphology 2: −2.37°, 4.77 Å). Resolutions were estimated by gold-standard FSC = 0.143 with the auto-tightened, noise-substituted mask: 2.71 Å and 2.91 Å (loose mask 3.01 and 3.18 Å). Local resolution was estimated with CryoSPARC Local Resolution Estimation (Supplementary Fig. 2). Helical symmetry was searched on a fixed 128 × 128 grid of twist and rise and imposed in real space over four asymmetric units once the refinement reached 8 Å, with the out-of-plane tilt limited to 10° and a dynamic mask (start 12 Å, near 6 Å, far 14 Å, z-clip 0.85). The Morphology 1 map was sharpened with B = −2.7 Å²; no sharpening was applied to the Morphology 2 map. The hand of the helix was not determined experimentally and remains unknown.

### Model building and refinement

Both maps show β-strands separated along the helical axis and clear side-chain density throughout the core, so for each morphology one cross-β layer was built de novo using ModelAngelo^24^ and Coot 0.9.8.96 ^25^ into the sharpened map and expanded to ten layers with the refined helical operator. Models were refined in Phenix 2.2.1 ^26^. Because Phenix places rotamers without regard to the identical copies of each residue in neighbouring layers, side chains were then re-optimized against the full ten-layer stack, scored with MolProbity and checked for loss of density fit. In Morphology 1, the Ser51–Tyr52 peptide of PF2 was flipped and Lys53 of the accessory chain, which has weak density (main chain 7.1 σ against a chain mean of 12.1 σ), was omitted. The final ten-layer Morphology 1 model was regenerated from its central layer with the refined helical operator. Atomic B-factors were refined last (real_space_refine run=adp, NCS constrained). Models were validated with MolProbity ^27^, phenix.validation_cryoem, EMRinger ^28^ and the wwPDB validation server (Supplementary Table 1). Solvent-accessible surface areas quoted in the text were computed from the final models with the Shrake–Rupley algorithm ^29^ (1.4 Å probe, 100 points per atom), buried area being the sum of the isolated-chain areas minus that of the complex. Medin numbering follows medin 1 = lactadherin 268 (UniProt Q08431); the deposited models 34WG and 34WH use lactadherin numbering 298–320 throughout. Structural figures were prepared in UCSF ChimeraX 1.12 ^30^; cross-sections of different structures shown side by side were rendered with one orthographic field width so that they are directly comparable.

### Aggregation propensity prediction

Aggregation propensity across medin (residues 1–50) was predicted with two complementary tools (Fig. 1a). WALTZ ^16^, a position-specific scoring matrix trained on experimentally characterized amyloid-forming hexapeptides, was run on the web server at pH 7.0 and segments were called at the high-sensitivity threshold of 79; at the default threshold only the N-terminal segment is called, so the choice is stated here. CORDAX ^17^, which scores the structural compatibility of a sequence with an amyloid core from high-resolution amyloid structures, was taken from the precomputed medin entry of the CORDAX server at its standard threshold, which covers medin 1–49. Both profiles and their provenance are available from the corresponding authors on request.

### FoldX energetics

Per-residue stabilities were computed with FoldX 5.1 ^31^. Each ten-layer deposition model (Morphology 1, three chains per layer; Morphology 2, four) was first processed with RepairPDB, and per-residue total ΔG values were then obtained from SequenceDetail on the repaired model. Chains were assigned to protofilament stacks by their position within each layer, and the two terminal layers were discarded because their solvent-exposed faces are not representative of the fibril interior; the per-residue ΔG reported for a stack is the mean over the remaining eight layers. For the landscape comparison the per-residue profile of each stack was smoothed with a five-residue sliding window and z-scored within the stack, following the procedure used to build the Amyloid Explorer database ^32^, and the medin C-APR stacks were placed against the 1,185 protofilament stacks of its 607 amyloid structures (42 proteins); a structure contributes one stack per protofilament, so most contribute two and the 1,185 is the sum over entries. The stack mean quoted in the text and plotted in Supplementary Fig. 7a excludes the first and last modelled residue of each stack, whose exposed termini are not representative of the stack, and the same exclusion is applied to every Amyloid Explorer stack; percentiles are of that stack mean against that distribution. FoldX inputs, outputs and the analysis scripts are available from the corresponding authors on request. FoldX reports ΔG_contrib as a sum of physical terms, and these were analysed alongside the total: backbone and side-chain hydrogen bonding, van der Waals packing and clashes, polar and apolar solvation, electrostatics, and the side-chain and backbone entropy costs of ordering. Term values quoted in the text are means per residue over the same inner layers, grouped into core and accessory stacks. For the frustration map, the per-residue ΔG_contrib of each stack was written into the B-factor column of one cross-β layer and coloured on a scale symmetric about zero. For the comparison with the full-length medin fibril (Supplementary Fig. 6d), the deposited coordinates of that structure (PDB 9U02) were processed identically. Its deposited model contains five layers per protofilament, so the ten-layer assembly supplied with that entry was used instead, with chains ordered by layer, to give each residue the same layer context as in the C-APR models; hydrogens were removed first. The resulting per-residue energies reproduce the Amyloid Explorer entry for 9U02 to r = 0.99 (mean absolute difference 0.10 kcal mol⁻¹ per residue), and an independently rebuilt ten-layer stack, generated by propagating the layer transform measured from the deposited coordinates, gives the same values to r = 0.99.

### Statistics and reproducibility

ThT kinetics are from one plate, with 4 wells of the C-APR and 4 of full-length medin measured side by side at 10 µM; individual wells are shown together with their mean, and an independent repeat on a second plate gave similar results. Negative-stain TEM was repeated twice with similar fibril morphologies; representative micrographs are shown. Cryo-EM data were collected once from one grid of a single fibril preparation; representative micrographs and 2D class averages are shown (Fig. 2). No statistical methods were used to predetermine sample sizes.

## Data availability

The cryo-EM maps (primary map, half maps and mask) have been deposited in the Electron Microscopy Data Bank under accession codes EMD-100201 (Morphology 1) and EMD-100202 (Morphology 2), and the atomic models in the Protein Data Bank under accession codes 34WG and 34WH. The unsharpened map of Morphology 1 is deposited as an additional map; no sharpening was applied to Morphology 2. Raw data are available from the corresponding author on request. The ThT kinetics and FoldX energies underlying the figures are available from the corresponding authors on request.

## Code availability

The custom scripts used for the FoldX stack analysis are available from the corresponding authors on request.

## Author contributions

B.P. produced the peptides, performed the aggregation assays, negative-stain and cryo-EM experiments, processed the data, built and refined the models, carried out the energetic analysis and prepared the figures. F.R. and J.S. designed and supervised the study and acquired funding. B.P. wrote the manuscript with input from all authors. All authors read and approved the final manuscript.

## Acknowledgements

B.P. is supported by a Junior Postdoctoral Fellowship (12AWB24N) from the Research Foundation – Flanders (FWO). This research was funded by grants from the FWO (G0C3522N) and the National Institutes of Health (NIH, R01AG079234). The content is solely the responsibility of the authors and does not necessarily represent the official views of the National Institutes of Health. Cryo-EM data were collected at the electron Bio-Imaging Centre (eBIC), Diamond Light Source, UK, proposal number BI41018, and we thank the eBIC staff for their support. We also thank Valentina Zorzini of the Biophysics Technology Unit, VIB-KU Leuven Center for Neuroscience. Finally, we would like to thank Rajasekhar Kolla, Ramon Duran Romana and Filip Claes for their review and suggestions in the drafting of the manuscript.

