## Supplementary figures and tables for "Two polymorphs of an independent amyloid core in medin’s C-terminal region"

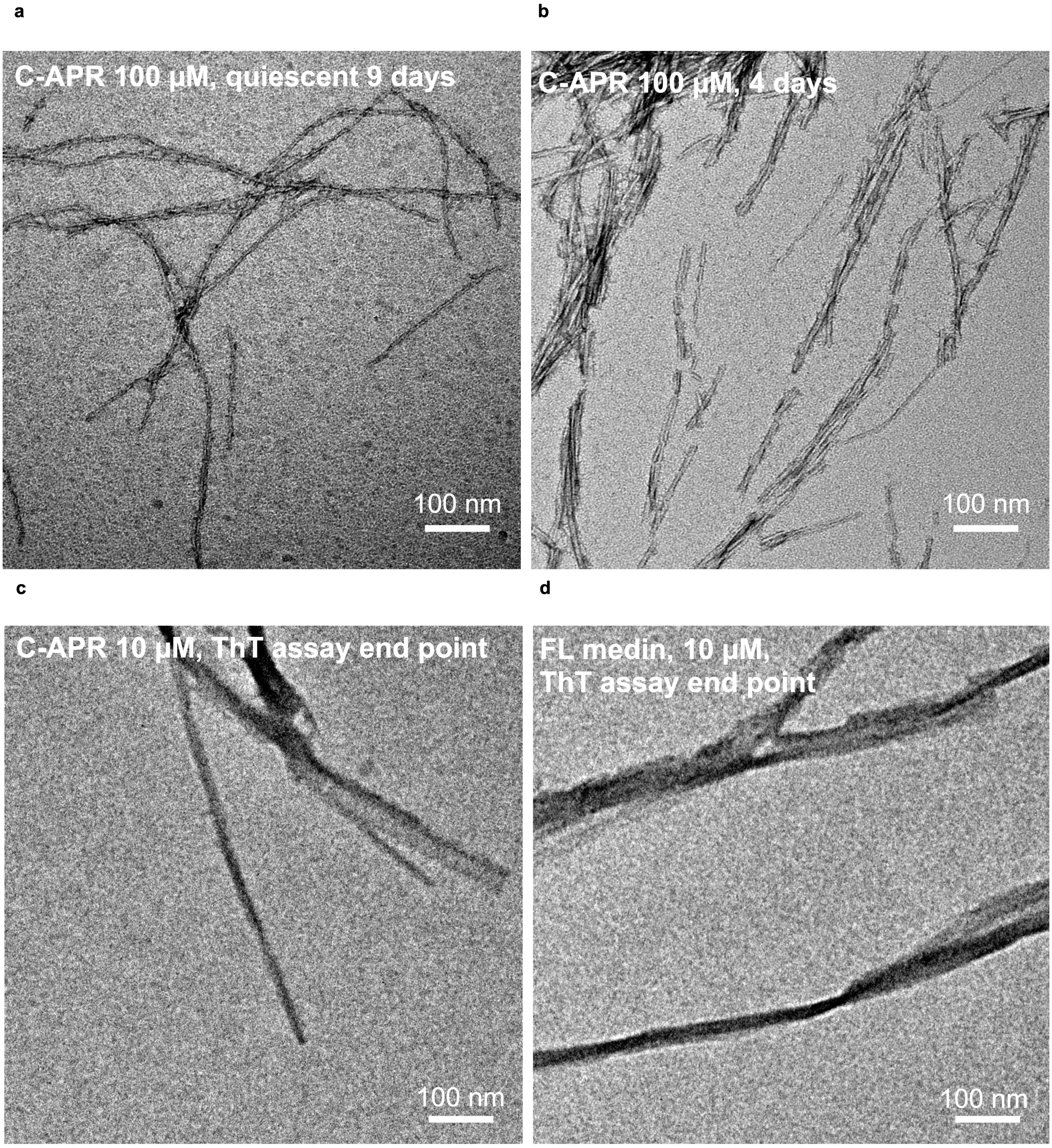


Supplementary Fig. 1 | The medin C-APR gives different fibril morphologies under different growth conditions. a, b, C-APR grown quiescently at 100 µM without dye, in two independent preparations four months apart: 9 days (a; the preparation used for cryo-EM) and 4 days (b). c, d, The end point of the thioflavin-T kinetic assay, 10 µM with a brief shake before each read: the C-APR (c) and full-length (FL) medin (d), from the same session and under the same conditions. All panels are 800 nm square fields; scale bars, 100 nm.

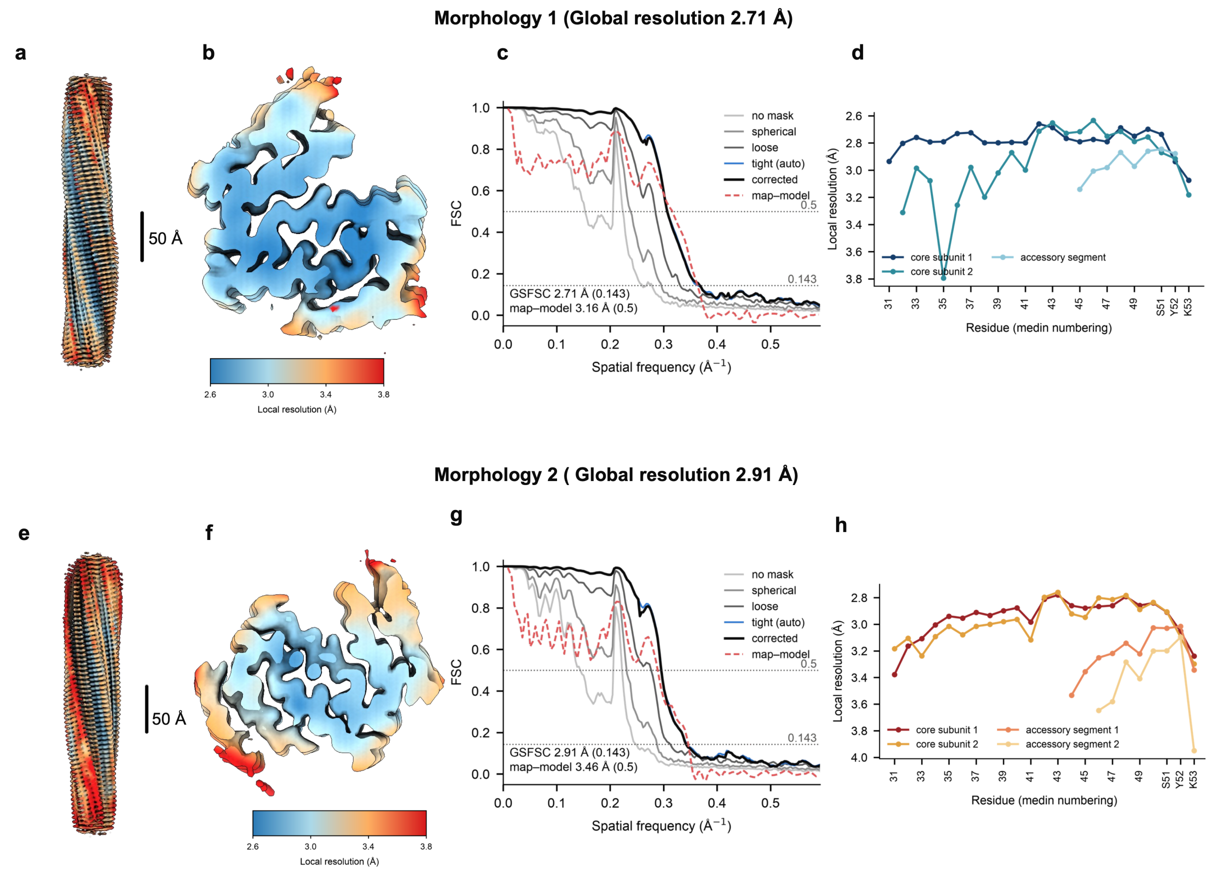


Supplementary Fig. 2 | Resolution assessment of the two reconstructions. a, e, Sharpened maps of Morphology 1 (a) and Morphology 2 (e) coloured by local resolution (CryoSPARC Local Resolution Estimation); scale bars, 50 Å. b, f, The same maps viewed along the fibril axis (12 Å slab). c, g, Gold-standard Fourier shell correlation between the two half maps with no mask, and with spherical, loose and auto-tightened masks, corrected by noise substitution; the 0.143 criterion gives 2.71 Å (c) and 2.91 Å (g). The red dashed curve is the map–model FSC (masked), crossing 0.5 at 3.16 Å (c) and 3.46 Å (g). d, h, Local resolution averaged over the eight inner layers of the deposited models, per residue and per chain. Residues are in medin numbering.

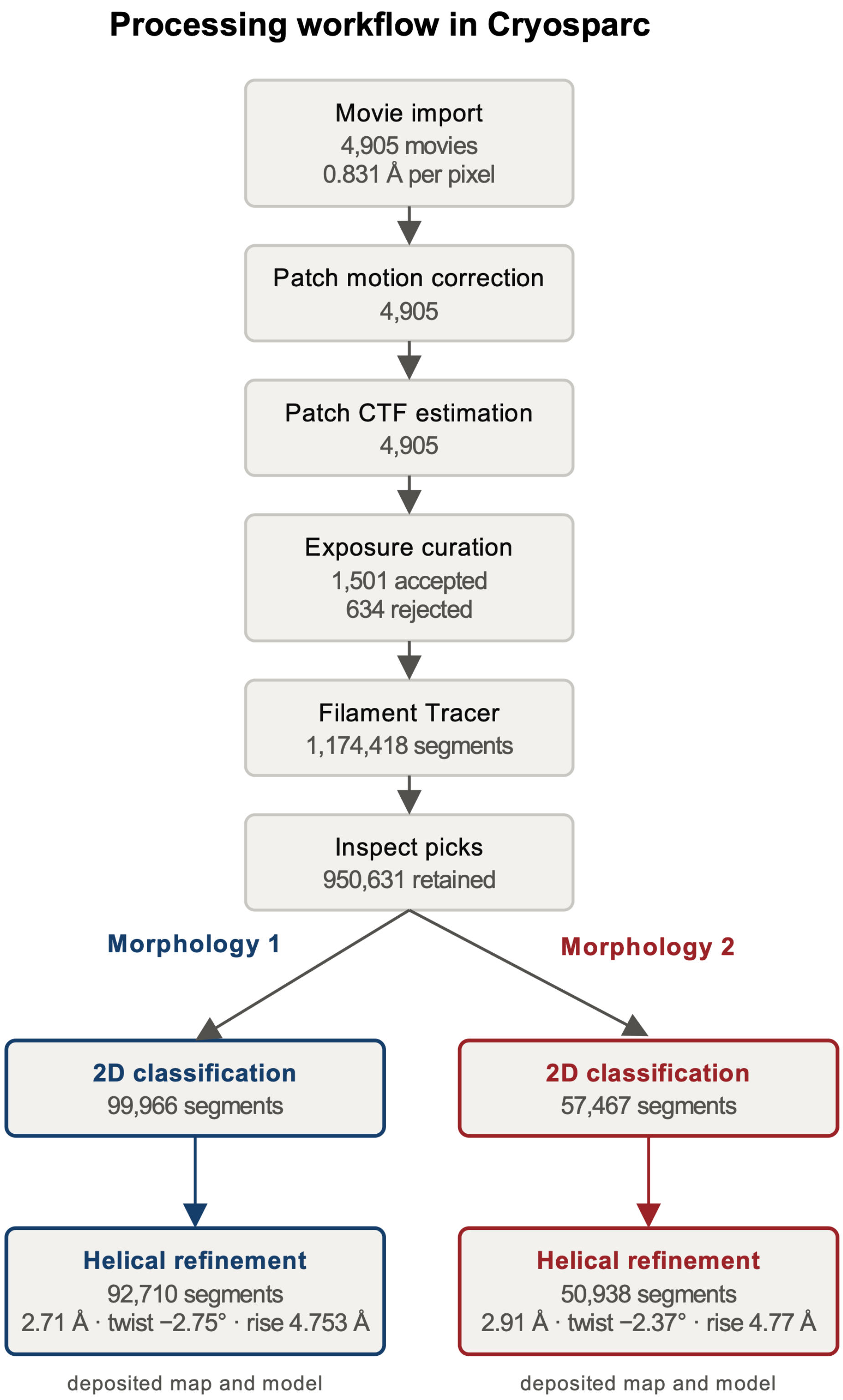


Supplementary Fig. 3 | CryoSPARC processing workflow. Workflow of project P68 with the number of items retained at each step; the two morphologies were separated by 2D classification and refined independently.

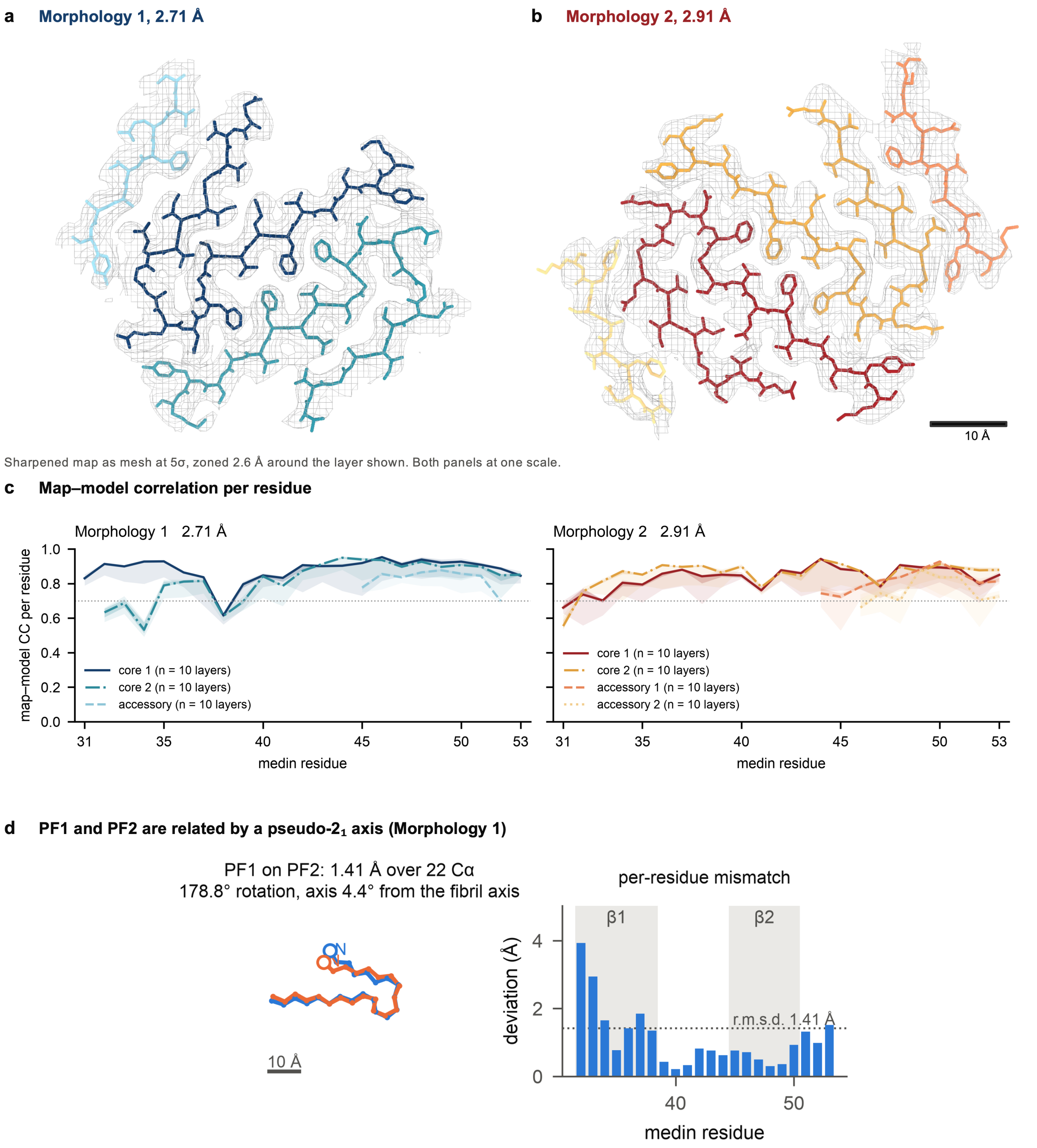


Supplementary Fig. 4 | Model geometry and map–model fit. a, b, Central layer of each deposited model shown as sticks inside the sharpened map, displayed as mesh at 5σ and zoned to 2.6 Å around the chains shown; cross-section perpendicular to the fibril axis, both panels at identical scale. Morphology 1 is shown in its cool palette (PF1 dark blue, PF2 teal, accessory segment pale blue) and Morphology 2 in its warm one (PF1 dark red, PF2 gold, accessory segments salmon and pale gold), as in Figs. 3 and 4. Scale bar, 10 Å. c, Map–model correlation per residue, in medin numbering; the line is the median over the ten layers of the deposited model, the dark band the interquartile range and the pale band the full range across layers. Median correlation per residue is 0.86 for Morphology 1 and 0.84 for Morphology 2, with 0.4 % and 0 % of residues below 0.5; overall CC_mask is 0.85 and 0.79 (Supplementary Table 1). d, The two core protofilaments of Morphology 1 are related by an approximate two-fold screw axis. Left, Cα traces of PF1 (dark blue) and PF2 (teal) after least-squares superposition (1.41 Å over 22 Cα, 178.8° rotation about an axis 4.4° from the fibril axis). Right, per-residue deviation after that superposition; shading marks β1 and β2, and the mismatch is confined to the N-terminal half of β1.

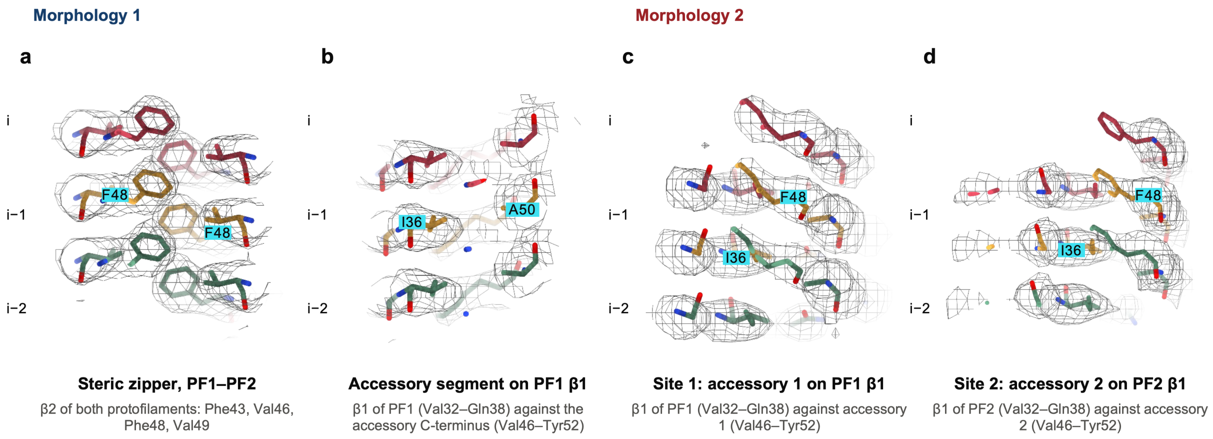


Supplementary Fig. 5 | Lateral views of the four interfaces. a, b, Morphology 1: the Phe43/Val46/Phe48/Val49 zipper between PF1 and PF2 (a) and the accessory segment against the β1 face of PF1 (b). c, d, Morphology 2: accessory segment 1 on the β1 face of PF1 (c) and accessory segment 2 on the β1 face of PF2 (d). Each panel shows three consecutive cross-β layers (i, i−1, i−2) coloured by layer, viewed perpendicular to the fibril axis, in the sharpened map (mesh). The transverse views of the same four interfaces are in Figs. 3e and 4e.

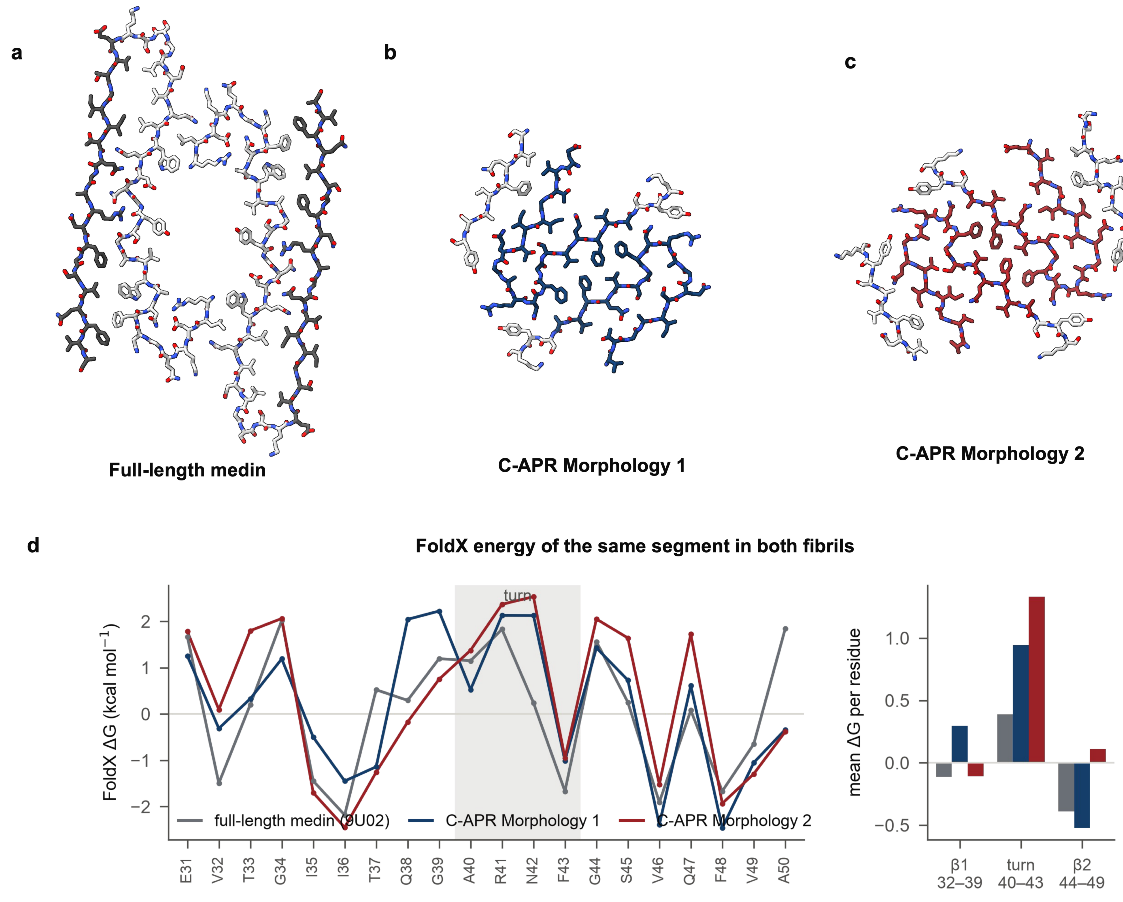

Supplementary Fig. 6 | The isolated medin C-APR adopts a fold unrelated to the same segment in the full-length medin fibril. a–c, Cross-sections perpendicular to the fibril axis, all three at identical scale: a, the full-length medin fibril (Pradhan, Kumar et al. 2026), one layer of its two protofilaments; b, C-APR Morphology 1; c, C-APR Morphology 2. The segment common to both constructs, medin 31–50 (lactadherin 298–317), is coloured; medin 1–30 in a and the accessory segments in b and c are light grey. Sequence identity over the shared segment was verified residue by residue. d, Per-residue FoldX ΔG over the same 20 residues: mean of the two protofilaments of the full-length fibril (grey) and of the two core chains of each C-APR polymorph (dark blue, Morphology 1; dark red, Morphology 2); shading, the Ala40–Phe43 turn. Right, segment means over medin 32–49. ΔG < 0 stabilizes the fold, ΔG > 0 marks a structurally frustrated residue. The two strands differ more between the polymorphs than either differs from the full-length fibril; the turn, by contrast, is two to three times as frustrated in both C-APR fibrils as in the full-length one.

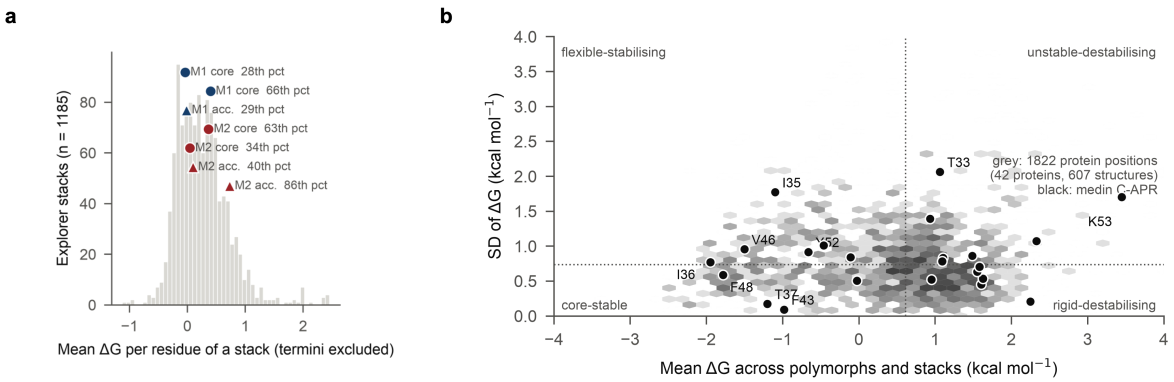


Supplementary Fig. 7 | The C-APR fibrils on the Amyloid Explorer energy landscape. a, Mean residue ΔG of each protofilament stack of the two polymorphs (circles, core; triangles, accessory) against all 1,185 Amyloid Explorer protofilament stacks (grey). The first and last modelled residue of every stack is excluded, here and in the database, and each label gives that stack's percentile of the distribution. b, Energy–variability landscape. For every protein position in the database (grey; 1,822 positions over 42 proteins and 607 structures) the mean ΔG across stacks is plotted against its standard deviation, with the 23 medin C-APR positions in black and the dotted lines at the database medians; the quadrant names are those of the Amyloid Explorer. ΔG < 0 stabilizes the fold, ΔG > 0 marks a structurally frustrated residue.

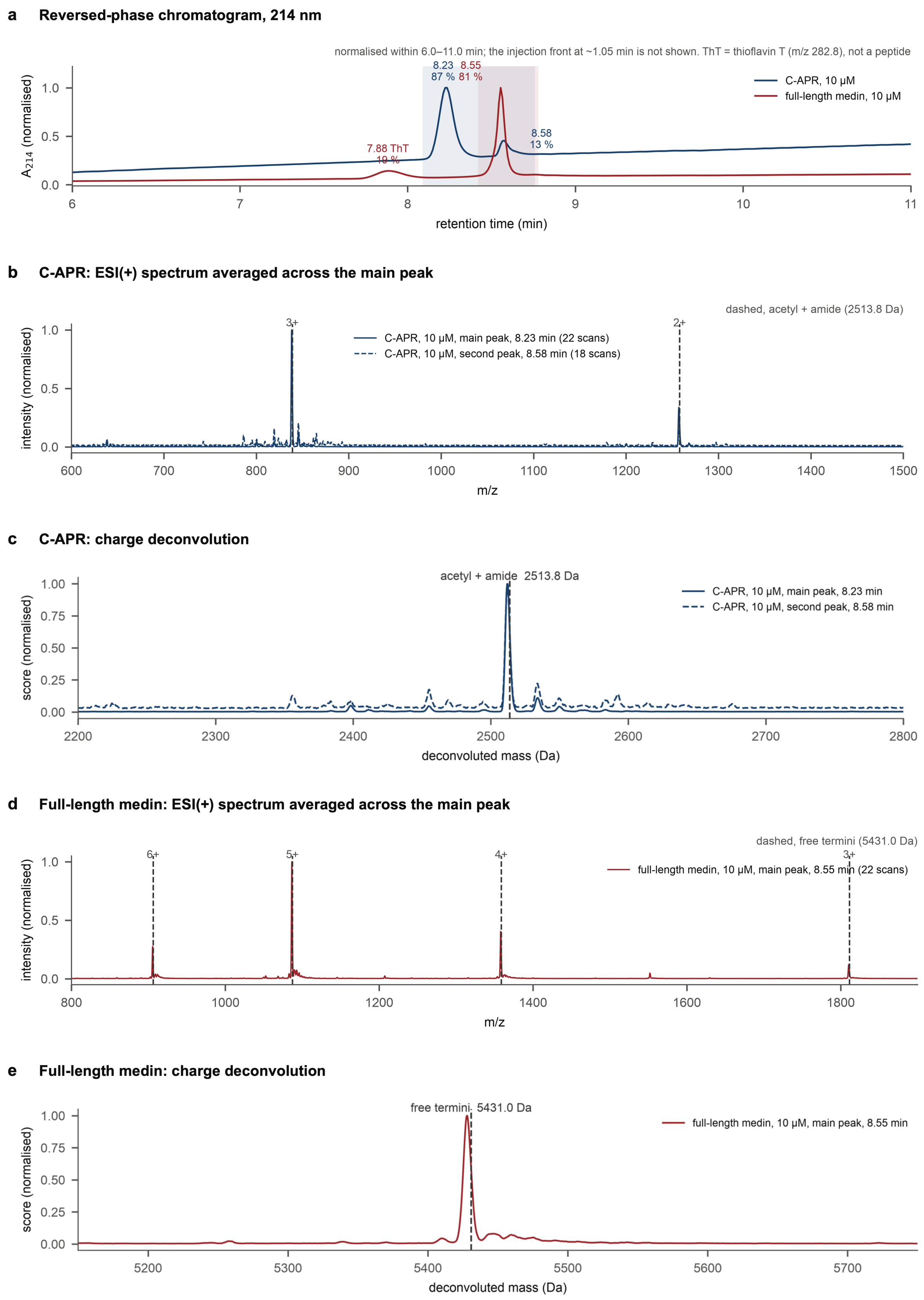


Supplementary Fig. 8 | LC–MS of the synthetic peptides. The C-APR and full-length medin, 10 μM each, injected back-to-back on the same day. a, Reversed-phase chromatograms (A214), normalized within the elution window; labels give each peak's retention time and its share of the integrated peak area, and shading marks the scans averaged for b and d. The

7.88 min peak of the medin run is thioflavin T, not a peptide: a single ion at m/z 282.8 with no charge ladder, matching the ThT cation ($C_{17}H_{19}N_2S^+$, 283.4 Da) once this run's calibration offset of −0.6 m/z is applied, with the M+2 isotope at 6.6 % as one sulfur predicts. b, ESI(+) spectra of the C-APR averaged across each of its two peaks (solid trace, 8.23 min; dashed trace, 8.58 min). Vertical dashed lines mark the m/z expected for the N-acetylated, C-amidated peptide at each charge state that carries an ion; the free-termini positions are not marked because no peak is observed there. c, Charge deconvolution of b. Both C-APR peaks give the same dominant species, 2,512.2 and 2,512.0 Da, assigned from independent charge states (2+ at m/z 1,257.2 and 1,257.1; 3+ at m/z 838.3), consistent with the capped peptide (2,513.8 Da); the free-termini mass (2,472.8 Da) accounts for 1 % of the base peak or less. The two C-APR peaks therefore differ in retention but not in mass. d,e, The same for full-length medin, where the marked positions are instead those of the free-termini peptide: it gives 5,428.0 Da, consistent with free termini (5,431.0 Da) and 44 Da below the capped mass (5,472.0 Da), and the capped positions, which are not marked, carry 0.7–6.8 % of the base peak. The two peptides therefore differ in their termini: the C-APR is capped, full-length medin is not.

Supplementary Table 1 | Cryo-EM data collection, refinement and validation statistics. (see Table1_cryoEM_statistics)

| | **Morphology 1 (J375)** | **Morphology 2 (J384)** |
|---|---|---|
| **Data collection and processing** | | |
| Microscope | Titan Krios | Titan Krios |
| Detector | Gatan K3 + BioQuantum | Gatan K3 + BioQuantum |
| Magnification | ×105,000 | ×105,000 |
| Voltage (kV) | 300 | 300 |
| Electron exposure ($e^-/Å^2$) | 40 (40 frames) | 40 |
| Defocus range (µm) | -0.9 to -2.8 | -0.9 to -2.8 |
| Pixel size (Å) | 0.831 | 0.831 |
| Movies collected / used | 4,905 / 1,372 | 4,905 / 1,292 |
| Symmetry imposed | C1; helical | C1; helical |
| Helical rise (Å) / twist (°) | 4.753 / −2.75 | 4.77 / −2.37 |
| Crossover (Å, from twist) | 311 | 362 |
| Initial segment images (no.) | 950,631 | 950,631 |
| Final segment images (no.) | 92,710 (15,689 filaments) | 50,938 (11,121 filaments) |
| Inter-box distance (Å) / box (px) | 21 / 400 | 21 / 400 |
| Map resolution (Å), FSC 0.143 | 2.71 | 2.91 |
| Local resolution (Å): best / median / 5–95 % (CryoSPARC J385, J386; refinement mask) | 2.42 / 3.45 / 2.7–7.0 | 2.60 / 3.70 / 2.9–7.7 |
| Map sharpening B factor ($Å^2$) | −2.7 | none applied |
| **Refinement** | | |
| Initial model used | de novo | de novo |
| Model resolution (Å), FSC 0.5 (masked) | 3.16 | 3.46 |
| Model–map FSC 0.143 (Å, masked) | 2.78 | 2.94 |
| Half-map FSC 0.143, Phenix (Å) | 2.70 | 2.83 |
| CC_mask / CC_volume | 0.85 / 0.83 | 0.79 / 0.78 |
| Model composition: chains / residues / atoms | 30 / 530 / 3990 | 40 / 640 / 4900 |
| Ligands | 0 | 0 |
| B factors ($Å^2$) min / max / mean | 42.44 / 133.84 / 81.01 | 58.81 / 150.69 / 93.24 |
| R.m.s. deviations: bond lengths (Å) | 0.009 | 0.004 |
| R.m.s. deviations: bond angles (°) | 1.290 | 0.923 |
| **Validation** | | |
| MolProbity score | 1.93 | 1.98 |
| Clashscore | 5.20 | 5.13 |
| Poor rotamers (%) | 0.00 | 0.00 |

| | | |
|---|---|---|
| Cβ deviations (%) | 0.00 | 0.00 |
| Ramachandran favoured (%) | 85.32 | 82.32 |
| Ramachandran allowed (%) | 14.68 | 17.68 |
| Ramachandran disallowed (%) | 0.00 | 0.00 |
| Rama-Z (whole) | -2.68 (0.37) | -5.57 (0.25) |
| EMRinger score | 4.21 | 3.43 |
| **Deposition** | | |
| EMDB / PDB | EMD-100201 / 34WG | EMD-100202 / 34WH |

Notes

Ramachandran values are Phenix/MolProbity on all ten layers, no hydrogens. MolProbity scores are computed from the MolProbity formula with these values (local MolProbity gives the same within 0.01).
